# Protocol-dependent cardiomyocyte states determine disease modelling capacity of human iPSCs

**DOI:** 10.64898/2026.03.29.715135

**Authors:** Sophie Shen, Clarissa Tan, Yuanzhao Cao, Chris Siu Yeung Chow, Dalia Mizikovsky, Janice Reid, Steve Dingwall, Andrew Prowse, Yuliangzi Sun, Zhixuan Wu, Sumedha Negi, Shaine Chenxin Bao, Enakshi Sinniah, Woo Jun Shim, Qiongyi Zhao, Jordan Thorpe, Azadeh Zahabi, Amy Hanna, Trishia Cheng, Adam Hill, James Hudson, James J. H. Chong, Nathan J. Palpant

## Abstract

Human induced pluripotent stem cell–derived cardiomyocytes (iPSC-CMs) are widely used to model cardiovascular disease, yet numerous differentiation protocols generate cardiomyocytes with heterogeneous molecular and functional properties, complicating experimental design. Here we systematically compare sixteen commonly used cardiomyocyte differentiation protocols and characterize their resulting cell states using single-nucleus RNA sequencing, functional phenotyping and computational integration with human genetic data. Despite similar cardiomyocyte yields, protocols produced distinct transcriptional programs, subtype compositions and physiological properties. By integrating protocol-specific gene expression signatures with genome-wide association studies of cardiovascular traits, we identify cardiomyocyte states enriched for genetic architectures underlying specific diseases. These analyses accurately predict protocols most suitable for modelling particular disease contexts, including electrophysiological defects associated with Brugada syndrome and metabolic vulnerability relevant to myocardial infarction. Our results demonstrate that differentiation protocols encode biologically distinct cardiomyocyte states with differential disease relevance and establish a framework for aligning stem-cell differentiation strategies with human complex trait genetics to guide model selection. This approach enables rational design of iPSC-based disease models and highlights how population-scale genetic data can inform experimental systems in stem cell biology.

## INTRODUCTION

Cardiovascular diseases (CVDs) remain the leading cause of death worldwide and encompass a diverse spectrum of disorders including inherited arrhythmias, cardiomyopathies and ischemic heart disease. Despite their scale and complexity, progress in therapeutic development has slowed, with cardiovascular drug pipelines experiencing persistent attrition and limited translation of preclinical discoveries into effective therapies.^1,2^ Across biomedical research, fewer than one in ten candidate drugs entering clinical trials ultimately succeed, highlighting the need for experimental systems that more faithfully capture human disease biology.^1,2^

A growing body of evidence suggests that limitations of conventional animal models contribute substantially to this translational gap. Cardiovascular physiology differs markedly across species, including differences in cardiac electrophysiology, ion channel expression and pharmacological responses that can complicate extrapolation of disease mechanisms and drug effects to humans.^3^ In response, governments and regulatory agencies have begun to actively promote the development of human-based experimental systems as alternatives to animal testing. The U.S. FDA Modernization Act 2.0, for example, allows drug developers to use human cell–based models and computational approaches in place of animal studies for regulatory evaluation, while initiatives such as the NIH Complement-ARIE program aim to accelerate adoption of next-generation human model systems across biomedical research. These policy shifts reflect a broader transition toward human-relevant experimental platforms capable of improving predictive power in drug discovery.

Human pluripotent stem cell (hPSC) technologies have emerged as one of the most promising approaches for achieving this goal. Human induced pluripotent stem cell–derived cardiomyocytes (iPSC-CMs) enable scalable generation of genetically defined human cardiac cells and are used to model disease mechanisms, evaluate pharmacological responses and study patient-specific genetic variation.^4,5^ However, the rapid expansion of this field has produced a large diversity of differentiation strategies that vary in developmental signaling logic, culture format and maturation approaches. A recent analysis of more than 300 published protocols highlighted substantial heterogeneity across differentiation methods and experimental readouts, raising a central methodological challenge for the field: how protocol choice shapes the resulting cardiomyocyte state and its suitability for disease modelling.^6^

Here we address this challenge by systematically comparing iPSC-cardiomyocyte differentiation protocols and characterizing the resulting cell states using single-nucleus transcriptomics and functional phenotyping. We further integrate protocol-specific transcriptional programs with genome-wide association studies of cardiovascular traits to assess the disease relevance of these states. Our results show that differentiation protocols generate distinct cardiomyocyte cell types with differential disease modelling capacity and establish a framework for aligning stem cell differentiation strategies with disease biology. This approach positions protocol diversity as a resource for designing more informative human models of cardiovascular disease.

## METHODS

### Generation and maintenance of human iPSC lines

All human pluripotent stem cell studies were carried out in accordance with consent from The University of Queensland’s Institutional Human Research Ethics approval (HREC#: 2015001434). Cardiomyocytes generated in this study were derived from the PGP1 hiPSC line. All PGP1 cells were maintained in mTeSR Plus medium with supplement (Stem Cell Technologies, Cat.#05825) at 37°C with 5% CO_2_. Cells were cultured on Vitronectin XF (Stem Cell Technologies, Cat.#07180) coated plates (Nunc, Cat.#150318).

### Cardiomyocyte differentiation protocols

Sixteen cardiomyocyte differentiation protocols were selected based on representation within the CMPortal protocol database^7^ and published literature describing human pluripotent stem cell differentiation strategies (Fig. 2a). Protocols were chosen to capture diversity in differentiation format, signalling modulation strategies and lineage specification approaches. Ventricular cardiomyocyte differentiation was performed using both three-dimensional embryoid body protocols and two-dimensional monolayer differentiation systems. Three-dimensional protocols (3D1 and 3D2) initiated differentiation through embryoid body formation followed by modulation of developmental signalling pathways. Two-dimensional protocols (2D1–2D4) used monolayer differentiation approaches with temporal activation and inhibition of Wnt signalling to induce cardiac mesoderm specification.

Wnt pathway activation was achieved using CHIR99021 and subsequently inhibited using small-molecule inhibitors including IWR2, IWP4 or XAV-939 depending on the protocol design. Variations in the dose, duration and timing of Wnt modulation were implemented according to published protocols and as shown in Fig 2a. Additional protocol variants were included to evaluate lineage specification and maturation interventions. Atrial cardiomyocyte differentiation was induced by introducing retinoic acid signalling during early cardiac mesoderm specification in modified versions of the 3D2 and 2D2 protocols (3D2+RA and 2D2+RA) and by using the STEMDiff Atrial Cardiomyocyte Differentiation Kit (STEMCELL Technologies). Ventricular enrichment was achieved in the 2D1+RAi protocol through inhibition of retinoic acid signalling during differentiation. A vascular-biased differentiation condition (2D3+VEGF) incorporated VEGF signalling to promote differentiation from multipotent mesoderm populations with endothelial potential. Two cardiomyocyte maturation conditions derived from the 2D4 protocol were included to evaluate post-specification maturation strategies. In the 2D4+RP+G condition, cardiomyocytes were replated during differentiation to induce hypertrophic and mechanotransductive signalling cues, whereas the 2D4+RP+FA condition incorporated fatty acid supplementation to promote metabolic maturation and oxidative metabolism. All differentiation protocols were maintained until day 15 prior to transcriptomic profiling.

### Flow cytometry

At the time of replating on day 15 of differentiation, a subset of cells (approximately 1 × 10^6^) was set aside for flow cytometry analysis of cardiomyocyte purity of differentiated cell populations. Cells were fixed with 4% paraformaldehyde (Sigma Aldrich, Cat.#158127-5G), permeabilised in 0.75% saponin (Sigma Aldrich, Cat.#S7900), and labelled with cardiac Troponin T antibody (ThermoFisher Australia Pty, Cat.# MA5-12960) or mouse IgG1 isotype control (Novus Biological, Australia Pty, Cat.# NBP1-43319). Stained samples were analysed using a LSRFortessa™ X-20 (Becton Dickinson) system with FACSDiva software (BD Biosciences). Data analysis was performed using FlowJo software 10 (version 10.6.2), and cardiac populations were determined with population gating from corresponding isotype controls. Cell preparations within 50% cardiac Troponin T-positive CMs were used.

### Single nuclei isolation

Nuclei isolation followed slight modification to 10X Genomic protocol which is described in detail below. Briefly, snap-frozen cell pellet was removed from ™ 80 °C storage and thawed on ice for 1min. To lyse the cells, 200uL cold Lysis Buffer were added into sample and dissociated with plastic pestle until homogenous. An additional 300uL cold lysis buffer was added to the remaining tissue sample and incubated on ice for 10min. Subsequently, sample was transferred into chilled nuclei isolation column in 2ml collection tube and centrifuged at 16,000 rcf for 20 secs in a 4 °C pre-cooled centrifuge. The nuclei isolation column was discarded and 2ml collection tube containing flow-through was vortexed at maximum speed for 10 sec to resuspend nuclei, then centrifuged at 500 rcf for 3 min in a 4 °C pre-cooled centrifuge. Supernatant was removed to 200uL in volume, leaving the pellet undisturbed. The nuclei pellet was resuspended in 500 µL chilled debris removal buffer and pipetted at least 15 times until no visible pellet observed, then centrifuged at 700 rcf for 10 min in a 4 °C pre-cooled centrifuge. Supernatant was removed to 200uL in volume, leaving the pellet undisturbed. The nuclei pellet was resuspended in 1 mL wash and resuspension buffer with 1:1000 Hoerscht. The 2mL tube containing nuclei suspension was placed in 50mL falcon tube cushioned with lint-free wipe and centrifuged at 500 rcf for 5 min in a 4 °C pre-cooled swing-bucket rotor centrifuge. The tube was carefully removed from 50mL falcon tube with a pair of forceps and supernatant was removed, leaving the pellet undisturbed. The nuclei pellet was resuspended in 260uL fixation buffer and pipetted at least 15 times until no visible pellet observed, then 10uL of nuclei sample was automatically counted using a Countess II FL automated cell counter (Thermo Fisher Scientific) following standard methodology. An additional 750uL fixation buffer was added to the remaining nuclei suspension and stored at 4°C for 16-24hours.

To sort nuclei, nuclei sample was removed from 4°C and placed in 50mL falcon tube cushioned with lint-free wipe and centrifuged at 850 rcf for 5 min in a room temperature swing-bucket rotor centrifuge. The tube was carefully removed from 50mL falcon tube with a pair of forceps and supernatant was removed, leaving the pellet undisturbed. The nuclei pellet was resuspended in 250uL cold quenching buffer and centrifuged at 850 rcf for 5 min in a room temperature swing-bucket rotor centrifuge. Supernatant was removed, leaving the pellet undisturbed and the nuclei pellet was resuspended in 250uL wash and resuspension buffer. Then a BD cell sorter was used to sort more than 1,000,000 Hoerscht positive events. The post-sort nuclei sample was centrifuged at 850 rcf for 5 min in a room temperature swing-bucket rotor centrifuge. Supernatant was removed to 50uL in volume, leaving the pellet undisturbed and nuclei pellet was resuspended in 250 uL quenching buffer, then 20uL of post-sort nuclei sample was evaluated under a microscope (EVOS, ThermoFisher) to ensure nuclei integrity and adequate count. The volume of post-sort nuclei was topped up to 1mL with quenching buffer with 100uL warmed enhancer and 275ul room temperature 50% glycerol and stored at ™ 80 °C for up to 3 months.

### snRNA-seq analysis

Gene expression matrices were normalized using log normalization and integrated across differentiation protocols using Seurat integration workflows. Principal component analysis was performed on the integrated dataset and dimensionality reduction was visualized using Uniform Manifold Approximation and Projection (UMAP). Graph-based clustering identified eleven major cellular clusters representing cardiomyocyte and non-cardiomyocyte populations. Cell type identities were assigned using a combination of automated annotation and manual evaluation. Automated classification was performed using scGPT cell type annotation models. Cluster identities were further validated through examination of canonical marker gene expression and enrichment analysis of Gene Ontology terms. Identified populations included cardiomyocytes, endothelial cells, fibroblasts, epithelial cells and endoderm-like cells. Cardiomyocytes were defined permissively as cells within clusters exhibiting strong cardiac muscle gene signatures or classified as cardiomyocytes by scGPT. To characterize cardiomyocyte subtype composition across differentiation protocols, label transfer analysis was performed using a published single-cell RNA sequencing reference dataset of human fetal heart development spanning 9–16 weeks post conception. Reference cardiomyocyte populations included early ventricular cardiomyocytes, late ventricular cardiomyocytes, atrial cardiomyocytes and proliferative cardiomyocytes. Subtype identities were assigned based on transcriptomic similarity to reference populations, and subtype proportions were quantified for each differentiation protocol. Transcription factor regulatory networks were inferred using the SCENIC framework. Gene co-expression networks were reconstructed from normalized expression matrices and transcription factor regulon activity scores were calculated for individual cells. Regulon enrichment across differentiation protocols was evaluated to identify protocol-specific regulatory programs. Metabolic pathway activity was inferred using the scFEA computational framework, which estimates metabolite flux states from single-cell transcriptomic data. Predicted metabolic fluxes across glycolysis, pyruvate metabolism and tricarboxylic acid cycle pathways were compared between differentiation conditions to assess metabolic maturation.

### Cardiomyocyte functional assays

Cardiomyocyte contractility was measured using high-resolution video microscopy. Beating cardiomyocytes were recorded and analyzed using motion-tracking software to quantify parameters including contractile amplitude, beat rate, relaxation velocity and upstroke velocity. To model myocardial infarction and ischemia–reperfusion injury, cardiomyocytes were subjected to hypoxic culture conditions followed by reoxygenation. Cell death was quantified using lactate dehydrogenase release assays. Electrophysiological properties were measured using multielectrode array recordings. Cardiomyocytes were plated onto MEA plates and spontaneous electrical activity was recorded to measure parameters including spike slope and depolarization rate.

### Integration with genome-wide association studies

To evaluate the disease relevance of protocol-specific cardiomyocyte states, transcriptomic data were integrated with genome-wide association study datasets. Differentially expressed genes for each protocol were identified relative to all other protocols. These gene sets were analyzed using SBayesRC to estimate enrichment of genetic variants associated with cardiovascular traits. Gene-based association analysis was performed using mBAT to identify genes contributing to heritability signals within UK Biobank GWAS datasets. GWAS traits analyzed included myocardial infarction, cardiac electrophysiological traits and arrhythmia syndromes.

### Statistics and reproducibility

All statistical analyses were performed using R. Data are presented as mean ± s.d. unless otherwise indicated. Comparisons between groups were performed using two-tailed Student’s t-tests or Wilcoxon rank-sum tests as appropriate. Multiple testing correction was applied using the Benjamini–Hochberg method where applicable. The number of biological replicates and statistical tests used for each experiment are reported in the corresponding figure legends. No statistical methods were used to predetermine sample size. Experiments were not randomized and investigators were not blinded to group allocation.

### Data availability

Single-nucleus RNA sequencing datasets generated in this study have been deposited in the Gene Expression Omnibus (GEO) under accession number and will be available upon publication. All code used for data processing and analysis will be available upon publication.

## RESULTS

### Trends in published protocol features for modelling cardiovascular disease

To understand how differentiation protocols are currently used across cardiovascular disease modelling studies, we expanded a recently published protocol database from Ewoldt et al.^6^ through systematic annotation and encoding to enable unsupervised analysis of protocol variables and study characteristics.^7^ This effort resulted in CMPortal, a searchable platform designed to interrogate relationships between protocol parameters, experimental design choices and disease modelling outcomes. CMPortal allows users to query a target variable such as a disease modelling endpoint and retrieve associated protocol features, study characteristics, cell line usage and phenotypic readouts reported across published protocols (**Figure 1a, Figure S1a**).

**Figure 1.**
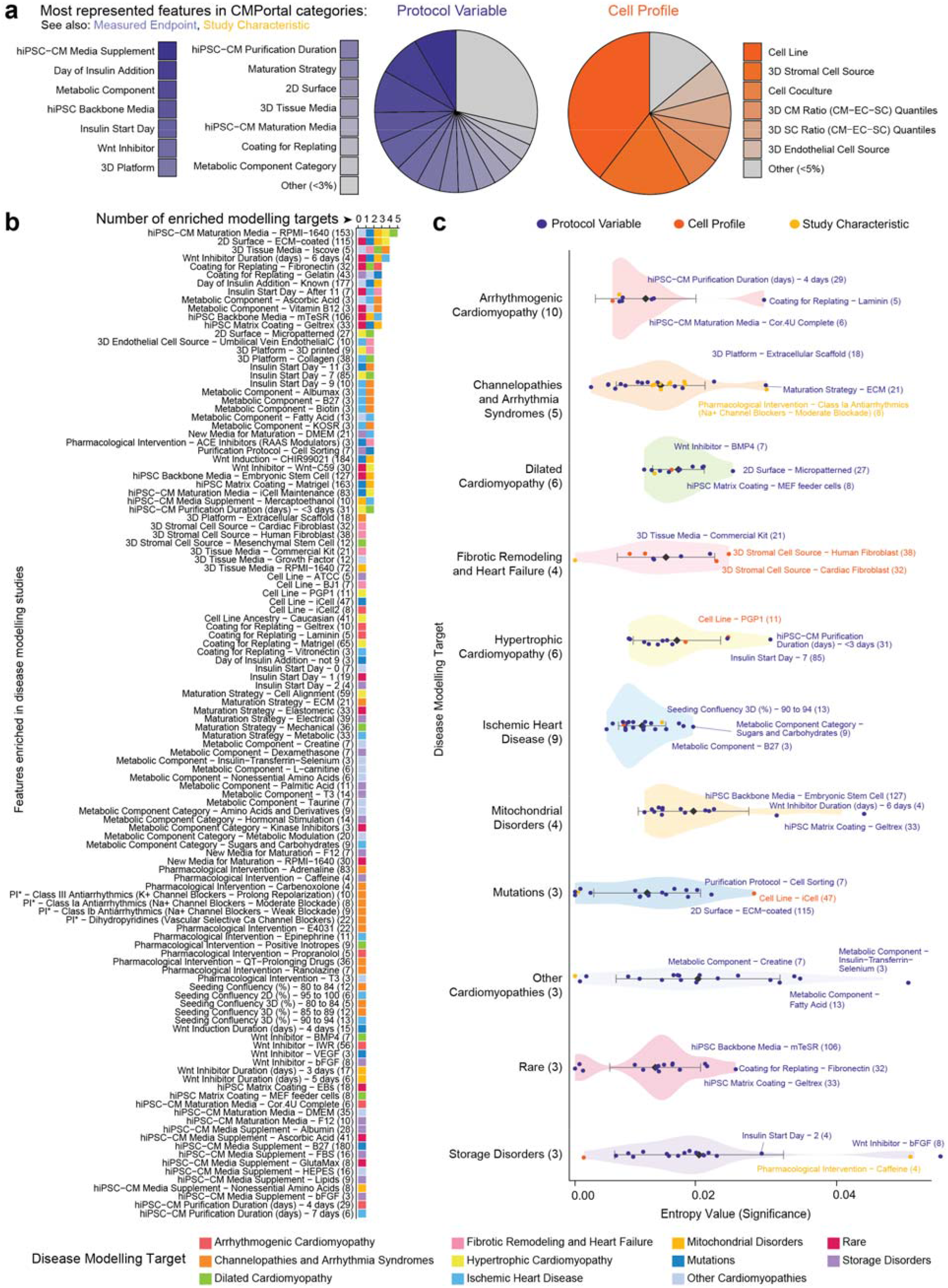
CMPortal identifies disease-specific patterns in published hiPSC-cardiomyocyte modelling studies. **a**, Overview of CMPortal categories used to annotate and query the published protocol corpus. Pie charts show the distribution of encoded features across protocol variable, cell profile, study characteristic and measured endpoint categories. CMPortal enables retrieval of associations between disease modelling targets and reported protocol parameters, cell line features, study design variables and phenotypic readouts. **b**, Number of enriched modelling targets for individual annotated features across the protocol corpus. Bars are colored by feature class (protocol variable, cell profile or study characteristic), and the histogram summarizes how many disease categories each feature is enriched in. Most enriched features were associated with a single disease modelling category, indicating low overlap in protocol design across disease contexts. **c**, Disease-specific enrichment of representative protocol features across cardiovascular disease modelling categories. Points indicate individual enriched features and are colored by feature class; horizontal position denotes entropy-based significance, with shaded violins showing the distribution within each disease category. Labels highlight the most prominent enriched variables for each target disease class, including extracellular matrix maturation and antiarrhythmic testing for channelopathies and arrhythmia syndromes, micropatterned culture conditions for dilated cardiomyopathy, fibroblast-containing 3D systems for fibrotic remodelling and heart failure, mid-differentiation insulin exposure for hypertrophic cardiomyopathy, metabolic components for ischaemic heart disease, embryonic stem cell media and Geltrex for mitochondrial disorders, and cell sorting or iCell lines for mutation-focused studies. Numbers in parentheses indicate the number of studies associated with each disease modelling category or annotated feature. See also Figure S1.

We used CMPortal to identify differentiation protocol features enriched within specific cardiovascular disease modelling categories. This analysis revealed striking divergence across disease contexts: of 118 enriched protocol features identified, 72.03% were associated with only a single disease category, indicating limited overlap in methodological approaches across disease modelling studies (**Figures 1b–c**). These patterns were consistent with expected biological associations. For example, studies modelling fibrotic remodelling following heart failure were enriched for cardiomyocyte–fibroblast co-culture systems (**Figure S1b**), whereas studies of arrhythmia syndromes frequently incorporated pharmacological testing of antiarrhythmic compounds (**Figure S1c**).

More broadly, disease-specific preferences were evident in differentiation and maturation strategies. Protocols used to study arrhythmias were more frequently associated with extracellular matrix–based maturation approaches, whereas studies of ischaemic heart disease more commonly incorporated metabolic maturation strategies (**Figure 1c, Figures S1c–d**). Additional trends included the use of mid-differentiation insulin exposure in hypertrophic cardiomyopathy models and the enrichment of micropatterned culture systems in studies of dilated cardiomyopathy (**Figures S1e–g**).

Together, these analyses indicate that protocol design is already implicitly tailored to specific cardiovascular disease contexts across the literature, but this heterogeneity has not been systematically evaluated or experimentally benchmarked. These observations motivated a controlled comparison of widely used cardiomyocyte differentiation protocols to determine how protocol-dependent cellular states influence their suitability for modelling distinct cardiovascular diseases.

### Selecting sixteen cardiomyocyte differentiation protocols for head-to-head comparison

To systematically evaluate how differentiation strategy influences cardiomyocyte state, we selected sixteen protocols spanning major methodological axes in the field, including culture format, Wnt signalling modulation, lineage biasing strategies and maturation approaches. The panel also included three commonly used commercial kits (PSC-CM: Gibco PSC Cardiomyocyte Differentiation Kit; SD-VCM and SD-ACM: STEMCELL Technologies STEMDiff Ventricular and Atrial Cardiomyocyte Differentiation kits) (**Figure 2a**).

**Figure 2.**
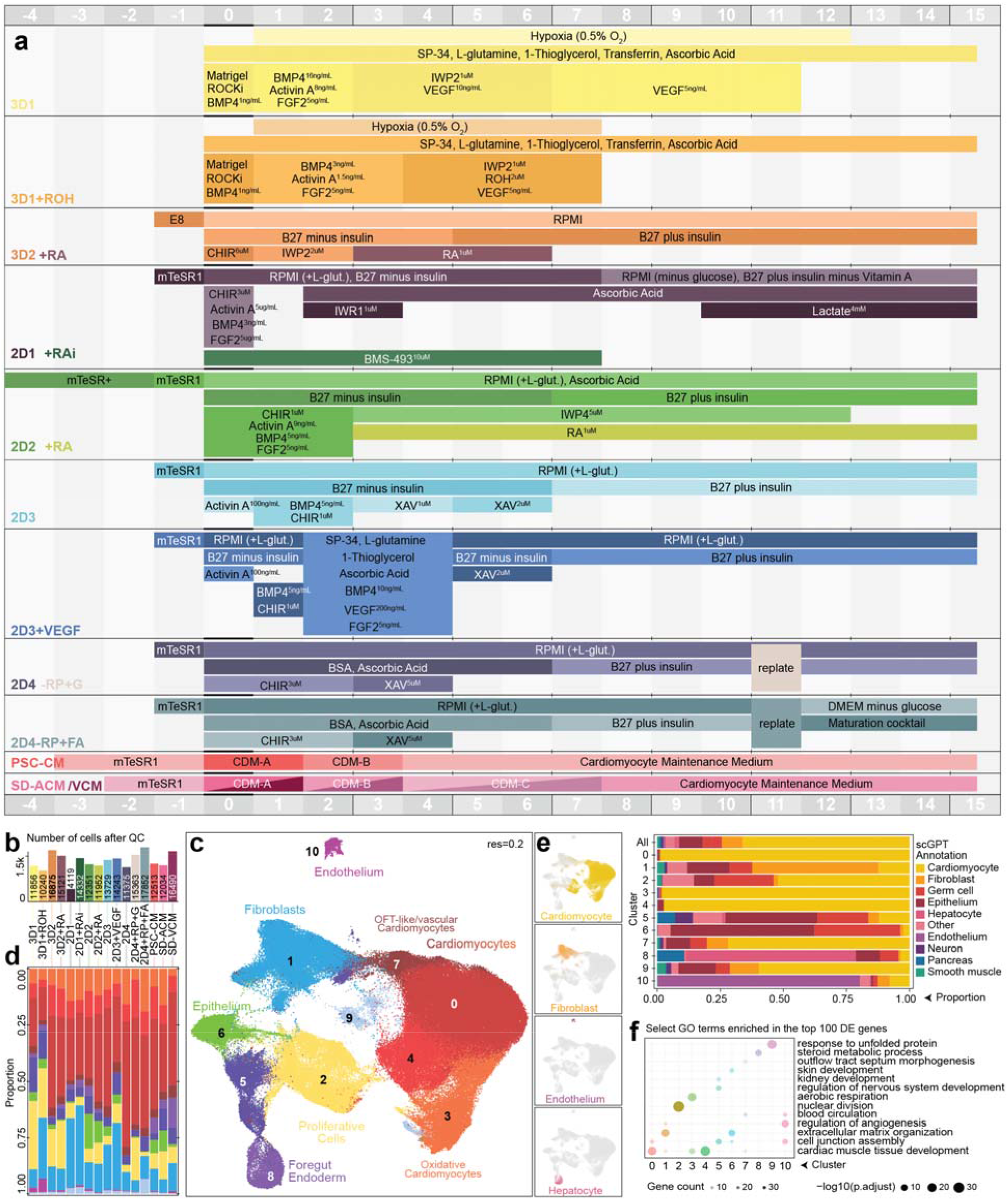
Differentiation protocol design and cellular composition of hiPSC-derived cardiac cultures. **a**, Schematic overview of the sixteen cardiomyocyte differentiation protocols compared in this study. Timelines indicate media composition, signalling modulators and lineage specification or maturation interventions from day ™4 to day 15 of differentiation. Protocols span major methodological axes used in the field, including three-dimensional embryoid body differentiation (3D1, 3D1+ROH, 3D2+RA), two-dimensional monolayer Wnt-modulation protocols (2D1+RAi, 2D2+RA, 2D3, 2D3+VEGF, 2D4 and derived maturation variants), and commercial differentiation kits (PSC-CM, SD-ACM and SD-VCM). Variations include modulation of Wnt activation and inhibition, retinoic acid– mediated atrial specification, VEGF-driven endothelial potential, and post-specification maturation strategies including replating-induced hypertrophy (2D4+RP+G) and fatty acid–based metabolic maturation (2D4+RP+FA). **b**, Number of high-quality nuclei recovered per protocol after quality control filtering of single-nucleus RNA sequencing datasets. **c**, UMAP visualization of integrated single-nucleus transcriptomes from day 15 differentiated cultures across all protocols. Graph-based clustering identified eleven transcriptionally distinct populations including cardiomyocytes, oxidative cardiomyocytes, OFT-like/vascular cardiomyocytes, proliferative cells, fibroblasts, endothelium, epithelial populations and foregut endoderm. **d**, Relative cellular composition across differentiation protocols. Stacked bar plots show the proportion of each annotated cell population within individual protocol-derived cultures following quality control. **e**, Cell type annotation of clusters using scGPT classification supported by marker gene expression. Stacked bars indicate the predicted identity distribution for each cluster across major cell classes including cardiomyocyte, fibroblast, endothelium, epithelium and other cell types. **f**, Representative Gene Ontology enrichment for genes highly expressed in each cluster. Dot plots display selected GO terms enriched among the top differentially expressed genes, with dot size representing gene count and colour indicating adjusted significance. These analyses support annotation of clusters corresponding to cardiomyocytes, proliferative populations and non-myocyte cell types within the differentiated cultures.

Five protocols represented ventricular cardiomyocyte differentiation across both three-dimensional embryoid body systems and two-dimensional monolayer formats. Two embryoid body protocols (3D1 and 3D2) differed primarily in their strategy for initiating cardiac specification. The 3D1 protocol uses a developmental mimicry approach with staged growth factor signalling, whereas 3D2 employs a simplified small-molecule strategy based on brief, high-intensity modulation of Wnt signalling. Four monolayer protocols (2D1–2D4) also varied in the timing and magnitude of Wnt pathway activation and inhibition. Protocol 2D1 uses a short high-dose CHIR99021 pulse followed by Axin stabilisation through IWR2-mediated inhibition of Wnt signalling. Protocol 2D2 employs lower-dose but prolonged CHIR exposure followed by sustained inhibition using IWP4. Protocol 2D3 similarly uses prolonged CHIR activation but suppresses Wnt signalling using XAV-939. In contrast, protocol 2D4 uses a short high-intensity CHIR pulse followed by brief inhibition with concentrated XAV-939. Unlike the other monolayer protocols, 2D4 excludes growth factor supplementation and relies exclusively on chemically defined small-molecule signalling modulation, reflecting a minimal differentiation strategy.^8,9^

To examine how lineage specification influences resulting cardiomyocyte states, several protocols were modified to bias differentiation toward distinct cardiac lineages. The 3D1 protocol was originally developed to produce first heart field–like cardiomyocytes, whereas an alternative variant (3D1+ROH) incorporates mesodermal signalling modulation to promote second heart field–like fates. Two ventricular protocols were additionally adapted to generate atrial cardiomyocytes through retinoic acid signalling (3D2+RA and 2D2+RA), which were compared with the proprietary STEMDiff atrial differentiation kit (SD-ACM). Conversely, the 2D1 protocol was modified to suppress atrial specification through inhibition of retinoic acid signalling (2D1+RAi), thereby enriching ventricular cardiomyocytes.

We also included protocol variants designed to probe alternative developmental trajectories and maturation states. The 2D3 protocol, originally developed to generate cardiomyocytes from multipotent mesoderm with endothelial potential,^10^ was supplemented with VEGF signalling (2D3+VEGF) to examine the resulting cellular composition. Finally, two maturation variants derived from the 2D4 protocol were incorporated to evaluate post-specification cardiomyocyte maturation strategies. These included replating-induced hypertrophic stimulation (2D4+RP+G) and metabolic maturation achieved through glucose deprivation and fatty acid supplementation (2D4+RP+FA).

Together, this panel of sixteen protocols captures the major methodological dimensions currently used to generate human iPSC-derived cardiomyocytes and provides a framework for directly comparing how differentiation strategy shapes cardiomyocyte identity and disease modelling capacity.

### Single-nuclei RNA sequencing of day 15 iPSC-derived cardiac cell types

To profile the cellular states generated by each differentiation strategy, we performed single-nucleus RNA sequencing (snRNA-seq) on day 15 differentiated cultures (**Figure 2a**). Across protocols, sequencing captured an average of 13,000 high-quality nuclei per sample (SD = 3,239.61), representing 75.42% of all captured nuclei after quality filtering (SD = 3.89%) (**Figure 2b**).

Unsupervised clustering of the integrated dataset identified 11 transcriptionally distinct cell populations, which were annotated using the scGPT classification framework supported by canonical marker gene expression and Gene Ontology (GO) enrichment analysis (**Figures 2c–f, Figure S2a–c**). Cardiomyocytes comprised majority of captured cells and were distributed primarily across clusters 0, 4, 3 and 7, with additional cardiomyocyte-like populations located at the periphery of clusters 2 and 9 (**Figure 2d**). Clusters 0 and 4 displayed the strongest enrichment for cardiac muscle gene programs, consistent with differentiated cardiomyocytes. Cluster 3 exhibited enrichment for oxidative metabolic pathways, indicating a more aerobic transcriptional profile, whereas cluster 7 showed mixed cardiomyocyte and vascular-associated signatures.

Cluster 2 was distinguished by strong expression of the proliferation marker MKI67 and enrichment for cell cycle– related transcripts, indicating a population of proliferative cardiomyocytes. In addition to cardiomyocytes, we detected several non-myocyte populations including vascular endothelial cells (cluster 10), fibroblasts (cluster 1), epithelial cells and endoderm-like populations (clusters 5 and 8).

Given that cardiomyocytes constituted the dominant cell population and were distributed across multiple transcriptional states, subsequent analyses focused on resolving cardiomyocyte subtype heterogeneity across differentiation protocols (**Figures S2a–b**).

### Characterisation of cardiomyocyte subtypes produced by different protocols

To compare cardiomyocyte identities produced by each differentiation strategy, we defined cardiomyocytes permissively as cells located in clusters 0, 4, 3, and 7, or classified as cardiomyocytes by scGPT (**Figures 3a–b**). Across protocols, this definition captured 27.32–86.73% of total cells, consistent with independent flow cytometry measurements of cardiac marker expression (**Figure S3a**).

**Figure 3.**
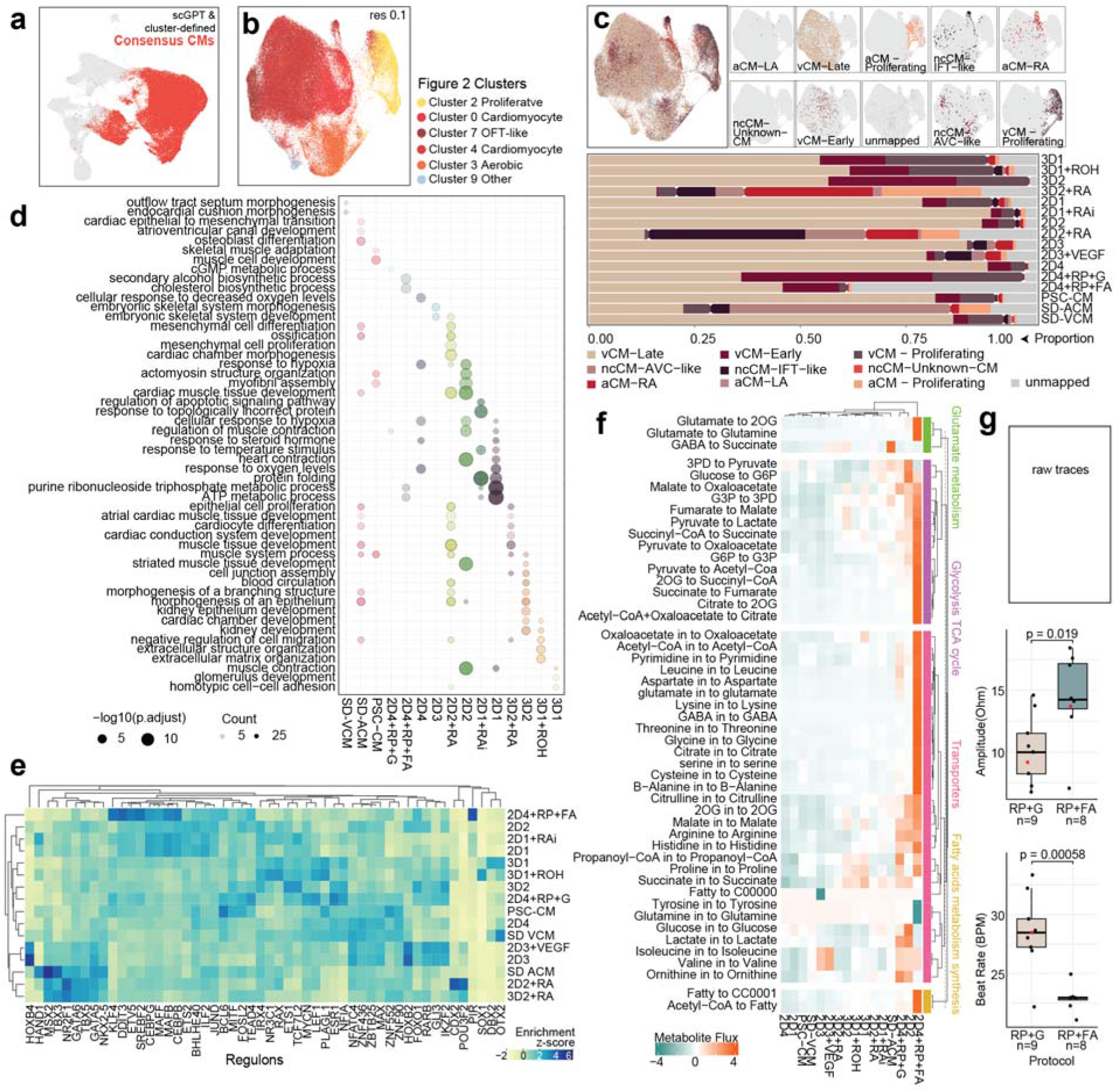
Cardiomyocyte subtype composition and maturation states across differentiation protocols. **a**, Identification of cardiomyocytes in the integrated single-nucleus dataset. UMAP projection showing consensus cardiomyocyte populations defined by agreement between scGPT cell type classification and cluster-based annotation. **b**, UMAP visualization of cluster identities derived from the integrated single-nucleus RNA-seq dataset (resolution = 0.1). Clusters correspond to cardiomyocytes, proliferative cells, OFT-like/vascular cardiomyocytes, oxidative cardiomyocytes and additional minor populations identified in Figure 2. **c**, Cardiomyocyte subtype classification based on label transfer to a human fetal heart single-cell reference atlas. UMAP projections show mapping to ventricular (vCM-Early and vCM-Late), atrial (aCM-RA and aCM-LA), atrioventricular canal–like (ncCM-AVC-like), inflow tract–like (ncCM-IFT-like), and proliferative cardiomyocyte states. The stacked bar plot summarizes the proportional representation of each cardiomyocyte subtype across differentiation protocols, revealing protocol-dependent enrichment of ventricular, atrial and proliferative cardiomyocyte states. **d**, Gene Ontology enrichment analysis of cluster-defining gene programs. Dot plots display selected enriched biological processes across clusters identified in the cardiomyocyte subset analysis. Dot size represents the number of genes associated with each GO term and colour indicates statistical significance (™log10 adjusted P value). **e**, Transcription factor regulatory network activity inferred using SCENIC. Heatmap showing regulon enrichment scores across differentiation protocols, highlighting protocol-specific regulatory programs associated with cardiomyocyte subtype identity and maturation state. **f**, Predicted metabolic pathway flux inferred using scFEA from single-nucleus transcriptomic data. Heatmap shows relative activity across metabolic reactions grouped into pathways including glutamate metabolism, glycolysis/TCA cycle activity, transport processes and fatty acid metabolism. The fatty acid maturation condition (2D4+RP+FA) shows increased flux through oxidative metabolic pathways relative to replated cardiomyocytes. **g**, Functional characterization of cardiomyocyte contractility comparing replated (2D4+RP+G) and fatty acid–matured (2D4+RP+FA) cardiomyocytes. Representative raw contraction traces are shown (top). Box plots quantify contractile amplitude and beat rate differences between conditions, demonstrating increased contractile strength and reduced spontaneous beating frequency following metabolic maturation. Statistical significance was assessed using two-tailed tests (P values shown).

To resolve cardiomyocyte subtype identity, we performed label transfer analysis using a reference single-cell RNA-seq atlas of human cardiac development spanning 9–16 weeks post conception.^11^ Mapping these reference populations onto our dataset defined three principal cardiomyocyte regions within the UMAP: ventricular cardiomyocytes (vCM-Early and vCM-Late), atrial or inflow tract–like cardiomyocytes (aCM-RA and ncCM-IFT-like), and proliferative cardiomyocytes of either ventricular or atrial identity (vCM-Proliferating and aCM-Proliferating), corresponding to the proliferative population observed in cluster 2 (**Figures 3b–c**).

Across differentiation strategies, the dominant subtype was late ventricular cardiomyocytes (vCM-Late). The highest proportions of this subtype were observed in monolayer ventricular differentiation protocols 2D2, 2D4, and the atrial lineage–inhibited 2D1+RAi condition (87.77%, 89.03%, and 89.78% respectively) (**Figure 3c**). The label transfer threshold was intentionally permissive (score ≥0.5), and the “late” ventricular reference population corresponds to mid-gestational cardiomyocytes rather than fully mature adult cells, as reflected by continued expression of immature sarcomeric isoforms (**Figure S3b**). Nevertheless, consistent differentiation trends emerged across protocols.

Three-dimensional differentiation protocols generated fewer vCM-Late cells and instead produced higher proportions of vCM-Early and proliferative cardiomyocytes, resembling the profile observed in replated monolayer cardiomyocytes (2D4+RP+G) (**Figure 3c**). In agreement with the whole-dataset clustering analysis (**Figure 2d**), the minimal small-molecule differentiation strategies 3D2 and 2D4 yielded the highest overall proportions of ventricular cardiomyocytes. In contrast, lineage-modifying variants of ventricular protocols (3D1+ROH and 2D3+VEGF) produced only modest changes in subtype distribution relative to their parent protocols. More pronounced shifts were observed in atrial differentiation and maturation conditions, indicating substantial remodeling of cardiomyocyte state in these contexts.

Atrial differentiation protocols (3D2+RA, 2D2+RA, and SD-ACM) exhibited markedly reduced proportions of vCM-Late cells and increased expression of atrial markers including NR2F2 and HCN4 (**Figures 3c–d, Figure S3c**). However, these protocols produced distinct atrial-like populations. The 3D2+RA protocol generated the largest proportion of right atrial and proliferative atrial cardiomyocytes (28.68% aCM-RA and 22.30% aCM-Proliferating). Transcription factor network analysis using SCENIC revealed enrichment of atrial-associated regulons TBX5 and TBX3, together with Gene Ontology terms related to cardiac conduction system development, consistent with a sinoatrial-like phenotype (**Figure 3d–e**).

In contrast, the 2D2+RA protocol produced a less specialized atrial myocardial population. Approximately 33.93% of cells mapped to atrial progenitor inflow tract–like cardiomyocytes (ncCM-IFT-like), accompanied by enrichment for gene programs related to chamber morphogenesis and circulation. Compared with 3D2+RA, this condition showed lower activation of the sinoatrial TBX3 regulon relative to the broader atrial TBX5 regulatory program.

The proprietary atrial differentiation kit (SD-ACM) generated a distinct cardiomyocyte state enriched for atrioventricular canal–like cells (ncCM-AVC-like; 49.33%), supported by enrichment of AVC-related GO terms and a TBX3^high^/TBX5^low^ regulon signature. This population also showed reduced expression of specialized atrial markers such as SHOX2 and NPPA and weaker alignment with atrial regions of the developing human heart (**Figure S3d**).

Finally, the 2D4 protocol family enabled comparison of two cardiomyocyte maturation strategies. Replating-induced maturation (2D4+RP+G) produced a shift from vCM-Late toward vCM-Early and proliferative cardiomyocyte states, consistent with hypertrophic growth. This phenotype was supported by enrichment of transcription factor regulons associated with cell growth and mechanotransduction, including TEAD4, LEF1, MYCN and ESR (**Figure 3c and e**).

In contrast, fatty acid supplementation (2D4+RP+FA) induced transcriptional programs associated with metabolic maturation. These included enrichment of sterol metabolism pathways (SREBF2 regulon and cholesterol biosynthetic processes), increased mitochondrial activity (MAFF and KLF4 regulons; ATP metabolic processes), and stress adaptation pathways (CEBPB, CEBPG and ETV5 regulons). Metabolic flux modelling using scFEA predicted increased pyruvate and tricarboxylic acid cycle flux, accompanied by reduced glucose and lactate uptake in fatty acid– treated cardiomyocytes (**Figure 3f**), consistent with the metabolic transition from glycolysis toward mitochondrial oxidative metabolism characteristic of cardiomyocyte maturation.

Functional measurements of contractility supported these transcriptional changes. Fatty acid–treated cardiomyocytes (2D4+RP+FA) exhibited increased contractile amplitude and reduced beat rate and relaxation velocity, features consistent with enhanced functional maturation relative to replated cardiomyocytes (2D4+RP+G) (**Figure 3g, Figure S3e**). An unexpected reduction in upstroke velocity was also observed after prolonged fatty acid treatment, suggesting a potential limitation in sustaining the energetic demands associated with more mature contractile activity (**Figure S3e–f**).

### Optimised protocols for specific disease modelling

The distinct cardiomyocyte states generated by different differentiation protocols suggested that protocol choice could influence the ability of iPSC-derived cardiomyocytes to model specific cardiovascular diseases. We therefore hypothesized that protocols enriched for transcriptional programs associated with particular disease traits would provide more appropriate in vitro models for those conditions. To test this hypothesis in an unbiased manner, we integrated cardiomyocyte transcriptomic profiles with human complex trait genetics. Using SBayesRC^12^, we estimated the contribution of genetic variants located in or near genes upregulated within each protocol-derived cardiomyocyte population to the heritable variance of multiple cardiovascular physiological and disease traits. This analysis revealed clear protocol-specific enrichments, indicating that even closely related differentiation strategies generate cardiomyocyte states aligned with distinct disease-relevant genetic architectures (**Figure 4a, Figure S4a–b**).

**Figure 4.**
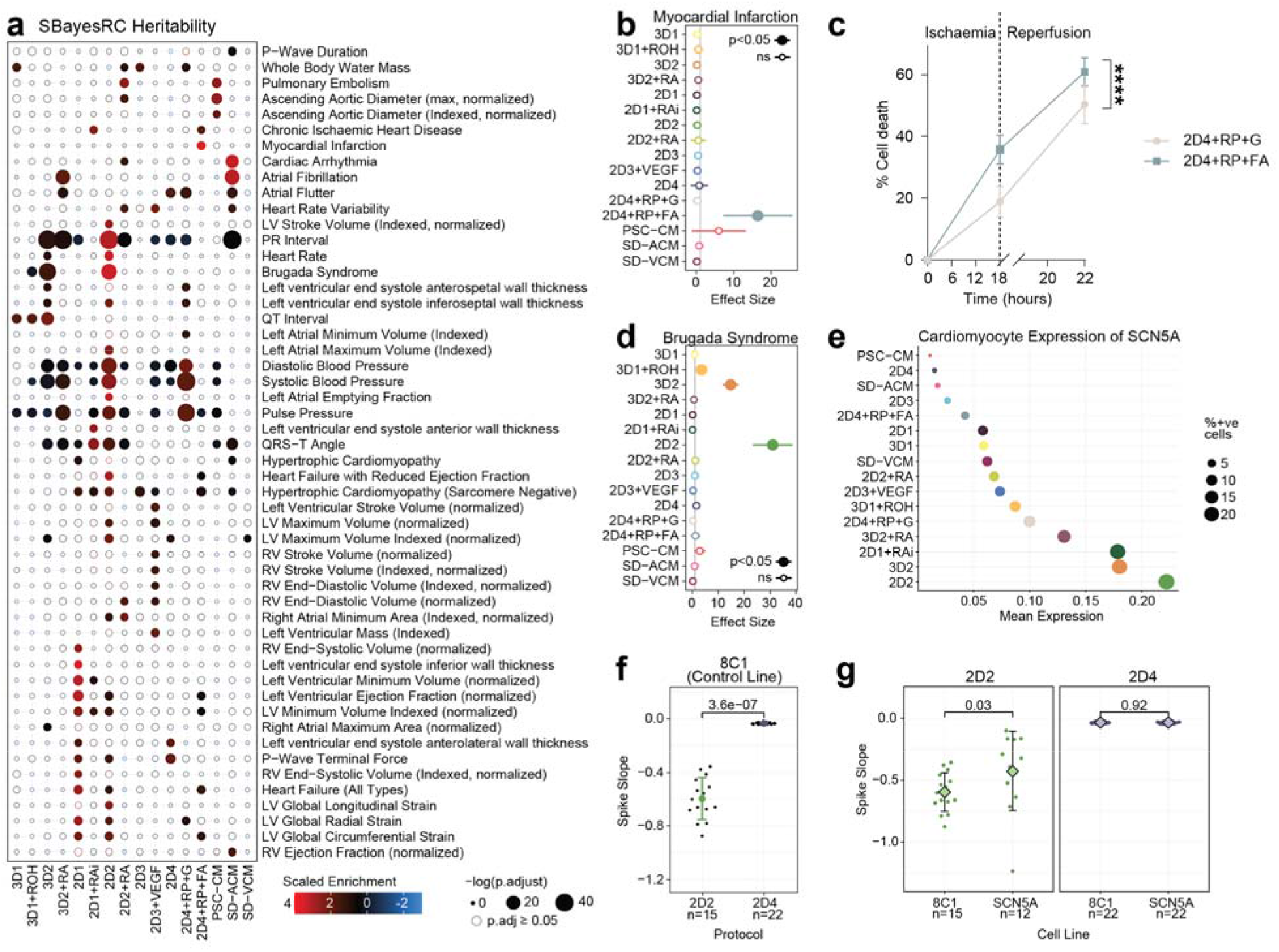
Integration of cardiomyocyte transcriptional programs with human genetic data predicts protocol suitability for disease modelling. **a**, Enrichment of cardiovascular trait heritability across protocol-derived cardiomyocyte states. Dot plot showing SBayesRC-derived scaled enrichment scores linking genes upregulated in each differentiation protocol to genome-wide association studies (GWAS) of cardiovascular traits. Rows represent traits and columns represent differentiation protocols. Dot colour indicates direction and magnitude of enrichment and dot size reflects statistical significance (™log10 adjusted P value). These analyses reveal protocol-specific enrichment of genetic architectures associated with cardiac electrophysiology, cardiac structure and cardiovascular disease traits. **b**, Predicted enrichment of myocardial infarction–associated genetic architecture across differentiation protocols based on gene–trait association analysis. Effect sizes represent enrichment of myocardial infarction GWAS signals within protocol-specific cardiomyocyte transcriptional programs. Statistical significance is indicated. **c**, Functional modelling of myocardial infarction and ischemia–reperfusion injury. Cardiomyocytes derived using replating-induced maturation (2D4+RP+G) or fatty acid–mediated metabolic maturation (2D4+RP+FA) were subjected to hypoxia followed by reperfusion. Line graph shows percentage of cell death over time quantified by lactate dehydrogenase release. Fatty acid–matured cardiomyocytes exhibit increased susceptibility to ischemic injury relative to replated cardiomyocytes. **d**, Predicted enrichment of Brugada syndrome genetic architecture across differentiation protocols. Effect sizes represent protocol-specific enrichment of GWAS signals associated with Brugada syndrome, highlighting ventricular protocol 2D2 as strongly enriched. **e**, Expression of the cardiac sodium channel gene SCN5A across cardiomyocytes generated by different differentiation protocols. Mean expression levels are plotted with point size indicating the proportion of cells expressing the gene, demonstrating highest expression in cardiomyocytes derived using protocol 2D2. **f**, Multielectrode array measurements of cardiomyocyte depolarization dynamics in a control iPSC line (8C1) differentiated using protocols 2D2 and 2D4. Spike slope values indicate stronger depolarization kinetics in 2D2-derived cardiomyocytes compared with 2D4-derived cells. **g**, Electrophysiological modelling of Brugada syndrome using an SCN5A mutant cell line. Spike slope measurements show reduced depolarization in SCN5A mutant cardiomyocytes relative to control cells when differentiated using protocol 2D2, whereas the phenotype is not detectable using protocol 2D4, demonstrating protocol-dependent sensitivity for modelling channelopathy phenotypes.

To functionally validate these predictions, we first examined two maturation variants derived from the 2D4 protocol: replating-induced hypertrophic stimulation (2D4+RP+G) and metabolic maturation via fatty acid supplementation (2D4+RP+FA). Genetic enrichment analysis predicted that the fatty acid–conditioned protocol would be more relevant for modelling myocardial infarction (MI) and ischemic injury (**Figure 4b, Figure S4a**). Gene-based mBAT analysis further supported this prediction, identifying enrichment of MI-associated variants near APOE in the fatty acid-treated condition (**Figure S4c**), consistent with the central role of lipid metabolism in cardiovascular disease. To experimentally test these predictions, we subjected cardiomyocytes to hypoxia followed by reoxygenation to model myocardial infarction and ischemia–reperfusion injury.^13^ Fatty acid–matured cardiomyocytes (2D4+RP+FA) exhibited significantly greater cell death compared with replated cardiomyocytes (2D4+RP+G), consistent with increased susceptibility to ischemic stress and supporting the prediction that this protocol better captures disease-relevant biology (**Figure 4c**).

We next assessed whether the framework could identify protocols optimized for modelling inherited cardiac channelopathies. Genetic enrichment analysis highlighted ventricular protocol 2D2 as strongly associated with Brugada syndrome, an inherited arrhythmia syndrome characterized by ventricular conduction abnormalities and increased risk of sudden cardiac death (**Figure 4d**). Although Brugada syndrome is classically associated with loss-of-function mutations in the cardiac sodium channel SCN5A, the disease is increasingly recognized as polygenic and mechanistically complex. Consistent with the genetic enrichment analysis, SCN5A expression was highest in cardiomyocytes derived from the 2D2 protocol, and mBAT analysis identified this locus as the dominant contributor to Brugada syndrome heritability signals in both 2D2 and 3D2 cardiomyocytes (**Figure 4e, Figure S4d**).

Functional electrophysiological measurements confirmed that protocol-dependent cardiomyocyte states influenced the ability to model disease phenotypes. Multielectrode array recordings showed that cardiomyocytes derived from protocol 2D4, which express relatively low levels of SCN5A, exhibited weaker depolarization dynamics compared with cardiomyocytes generated using 2D2 (**Figure 4f**). Importantly, the stronger depolarization kinetics of 2D2-derived cardiomyocytes enabled robust detection of reduced depolarization rates in an SCN5A mutant cell line, whereas the phenotype was not clearly detectable in the 2D4 background (**Figure 4g**).

## DISCUSSION

Selecting appropriate cellular models remains a fundamental challenge in stem-cell–based disease modelling. Differentiation protocols for iPSC-derived cardiomyocytes vary widely yet are often treated as interchangeable routes to generate a generic cardiomyocyte population. Here we introduce a framework that integrates transcriptomic profiling with human complex trait genetics to systematically align differentiation protocols with disease mechanisms.

By linking protocol-specific cardiomyocyte gene programs to GWAS-derived heritability across cardiovascular traits, we show that different differentiation strategies generate cardiomyocyte states with distinct disease relevance. This approach transforms protocol selection from empirical convention into a data-driven decision guided by the genetic architecture of the disease being studied.

Recent studies have begun to demonstrate how stem-cell differentiation systems can be used to interpret human genetic variation. We recently showed that transcriptional programs emerging during iPSC lineage differentiation could be integrated with GWAS signals to identify regulators of cardiovascular traits, leading to the discovery of *TMEM88* as a modulator of mammalian blood pressure^14^. Similarly, mapping *HOPX*-dependent regulatory networks in differentiating cardiomyocytes demonstrated that these developmental programs intersect with genetic architectures governing human cardiac traits^15^. Complementary work mapping eQTLs in iPSC-derived cardiomyocytes has further shown that many cardiovascular GWAS variants exert regulatory effects in specific cellular contexts. Collectively, these studies establish that stem-cell differentiation models can reveal biological mechanisms underlying complex traits.

Our results extend this concept in an important way. Rather than linking genetic loci to individual genes or regulatory networks, we show that differentiation protocols themselves define cellular contexts that differentially capture disease-associated genetic programs. By integrating cardiomyocyte transcriptomes from multiple protocols with GWAS data, we identify differentiation strategies enriched for genetic architecture relevant to specific cardiovascular diseases. Importantly, these predictions translated into improved functional disease modelling, as illustrated by identification of a ventricular protocol capable of resolving electrophysiological defects associated with Brugada syndrome. In this framework, human genetics can be used not only to interpret disease mechanisms but also to guide the selection of experimental systems most likely to reveal those mechanisms.

This perspective offers a potential explanation for a longstanding source of variability in iPSC disease models. If differentiation protocols generate cardiomyocytes with distinct regulatory states, then their ability to reveal disease phenotypes will depend on whether those states intersect with the biological pathways underlying the disease. Our results show that this alignment can be quantified and leveraged experimentally. Rather than searching for a universally optimal differentiation method, different protocols may provide complementary cellular contexts optimized for different classes of cardiovascular disease.

Several limitations should be noted. Our comparison of sixteen representative protocols captures common differentiation strategies but does not systematically resolve the causal contribution of individual signalling variables. The predictive framework also depends on currently available protocol annotations and GWAS datasets, both of which remain incomplete. Nevertheless, the strong concordance between computational predictions and functional disease modelling supports the utility of integrating transcriptomic and genetic data to guide model selection.

More broadly, this work illustrates how population-scale genetic data can inform the design of experimental systems in stem-cell biology. Integrating differentiation state space with complex trait genetics provides a general strategy for matching in-vitro cell models to disease mechanisms. Applying this approach across additional cell types and differentiation systems may enable more rational design of stem-cell models for studying human disease.

## REFERENCES

1 Roth, G. A. et al. Global Burden of Cardiovascular Diseases and Risk Factors, 1990–2019: Update From the GBD 2019 Study. Journal of the American College of Cardiology 76, 2982–3021 (2020).

2 Sun, D., Gao, W., Hu, H. & Zhou, S. Why 90% of clinical drug development fails and how to improve it? Acta Pharmaceutica Sinica B 12, 3049–3062 (2022).

3 Hamlin, R. L. & Keene, B. W. Species differences in cardiovascular physiology that affect pharmacology and toxicology. Current Opinion in Toxicology 23-24, 106–113 (2020).

4 Thomas, D., Cunningham, N. J., Shenoy, S. & Wu, J. C. Human-induced pluripotent stem cells in cardiovascular research: current approaches in cardiac differentiation, maturation strategies, and scalable production. Cardiovascular Research 118, 20–36 (2021).

5 Raniga, K. et al. Strengthening cardiac therapy pipelines using human pluripotent stem cell-derived cardiomyocytes. Cell Stem Cell 31, 292–311 (2024).

6 Ewoldt, J. K. et al. Induced pluripotent stem cell-derived cardiomyocyte in vitro models: benchmarking progress and ongoing challenges. Nature Methods 22, 24–40 (2025).

7 Chris Siu Yeung Chow, S.N., Shaine Chenxin Bao, Chen Fang, James E. Hudson, Woo Jun Shim, Yuanzhao Cao, Nathan J. Palpant. A community-oriented, data-driven resource to improve protocol design for cardiac modelling from human pluripotent stem cells. Biorxiv (2024).

8 Burridge, P. W. et al. Chemically defined generation of human cardiomyocytes. Nat Methods 11, 855–860 (2014).

9 Lian, X. et al. Robust cardiomyocyte differentiation from human pluripotent stem cells via temporal modulation of canonical Wnt signaling. Proc Natl Acad Sci U S A 109, E1848–1857 (2012).

10 Palpant, N. J. et al. Inhibition of beta-catenin signaling respecifies anterior-like endothelium into beating human cardiomyocytes. Development 142, 3198-3209 (2015).

11 Farah, E. N. et al. Spatially organized cellular communities form the developing human heart. Nature 627, 854-864 (2024).

12 Zheng, Z. et al. Leveraging functional genomic annotations and genome coverage to improve polygenic prediction of complex traits within and between ancestries. Nat Genet 56, 767-777 (2024).

13 Redd, M. A. et al. Therapeutic Inhibition of Acid-Sensing Ion Channel 1a Recovers Heart Function After Ischemia-Reperfusion Injury. Circulation 144, 947-960 (2021).

14 Shen, S. et al. Atlas of multilineage stem cell differentiation reveals TMEM88 as a developmental regulator of blood pressure. Nat Commun 16, 1356 (2025).

15 Friedman, C. E. et al. HOPX-associated molecular programs control cardiomyocyte cell states underpinning cardiac structure and function. Developmental Cell 59, 91-107.e106 (2024).

