## SUPPLEMENTAL INFORMATION for "Protocol-dependent cardiomyocyte states determine disease modelling capacity of human iPSCs"

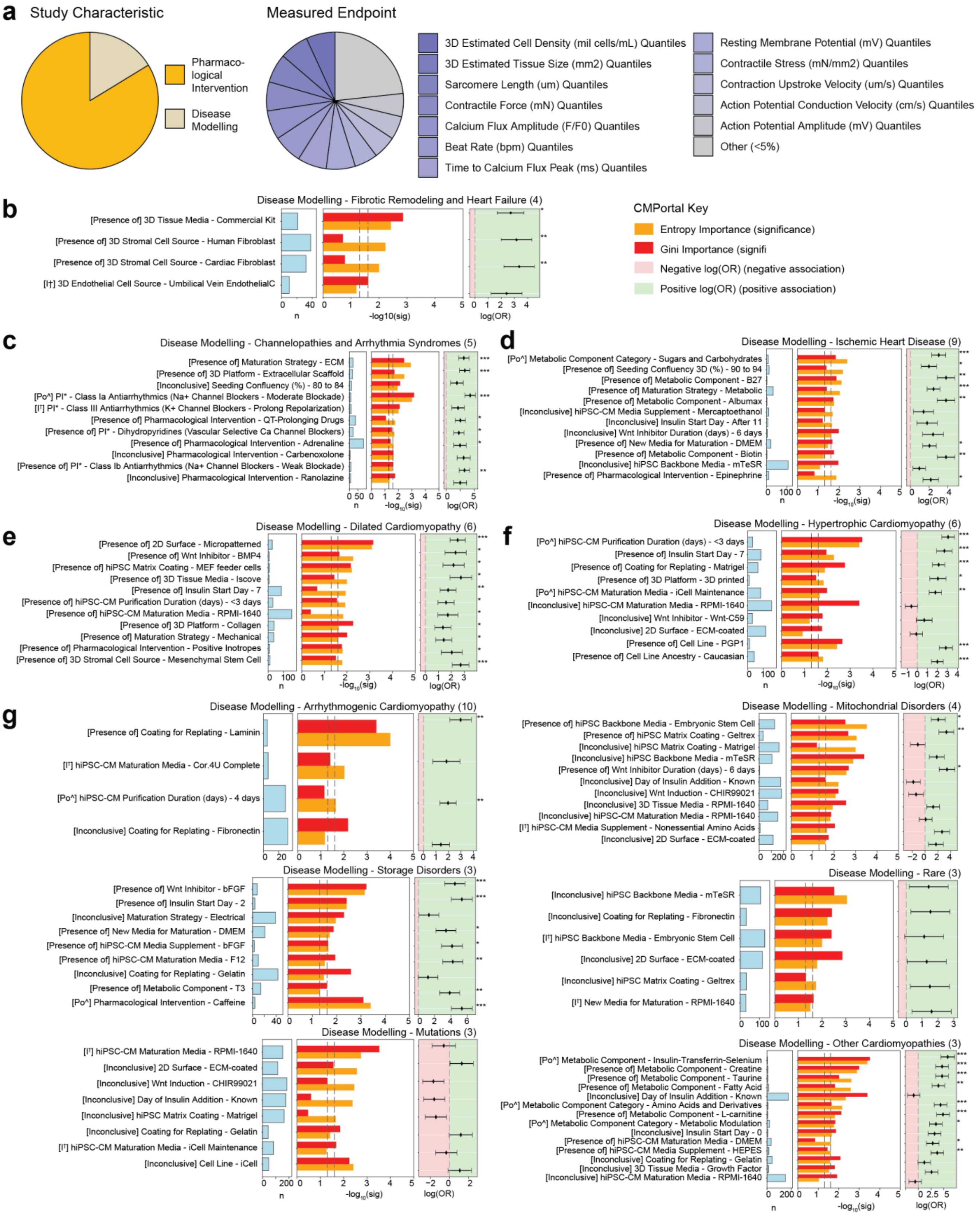

**Figure S1 | CMPortal analysis linking cardiomyocyte differentiation protocol features to disease modelling contexts.**

**a,** Distribution of study characteristics and measured endpoints across the CMPortal protocol database. Pie charts summarize the proportion of studies categorized as pharmacological intervention or disease modelling (left) and the diversity of functional readouts reported in published cardiomyocyte studies (right), including measurements of sarcomere length, contractile force, calcium flux amplitude, beat rate, action potential parameters and tissue-level structural properties.

**b–g,** CMPortal-derived enrichment analysis identifying protocol variables associated with specific cardiovascular disease modelling categories. For each disease context, the left panel shows the number of studies containing a given feature, the middle panel shows feature importance based on entropy (orange) and Gini (red) significance metrics, and the right panel shows effect size represented as log odds ratios with confidence intervals. Green shading indicates positive association and pink shading indicates negative association with the disease modelling category.

**b,** Protocol features enriched in studies modelling fibrotic remodelling and heart failure, including three-dimensional tissue systems and fibroblast-containing co-culture conditions.

**c,** Features associated with studies of channelopathies and arrhythmia syndromes, highlighting extracellular matrix maturation strategies, electrophysiological assays and pharmacological testing of antiarrhythmic compounds.

**d,** Protocol characteristics enriched in studies modelling ischaemic heart disease, including metabolic maturation strategies, specific metabolic media formulations and small-molecule signalling modulators.

**e,** Protocol features associated with dilated cardiomyopathy studies, including micropatterned culture substrates, mechanical maturation approaches and specific cardiomyocyte purification or maturation conditions.

**f,** Protocol variables enriched in hypertrophic cardiomyopathy studies, including insulin supplementation timing, three-dimensional culture platforms and specific cardiomyocyte maturation media.

**g,** Additional disease modelling categories including arrhythmogenic cardiomyopathy, mitochondrial disorders, storage disorders, rare diseases and other cardiomyopathies. These analyses highlight disease-specific enrichment of protocol design parameters across the published cardiomyocyte differentiation literature.

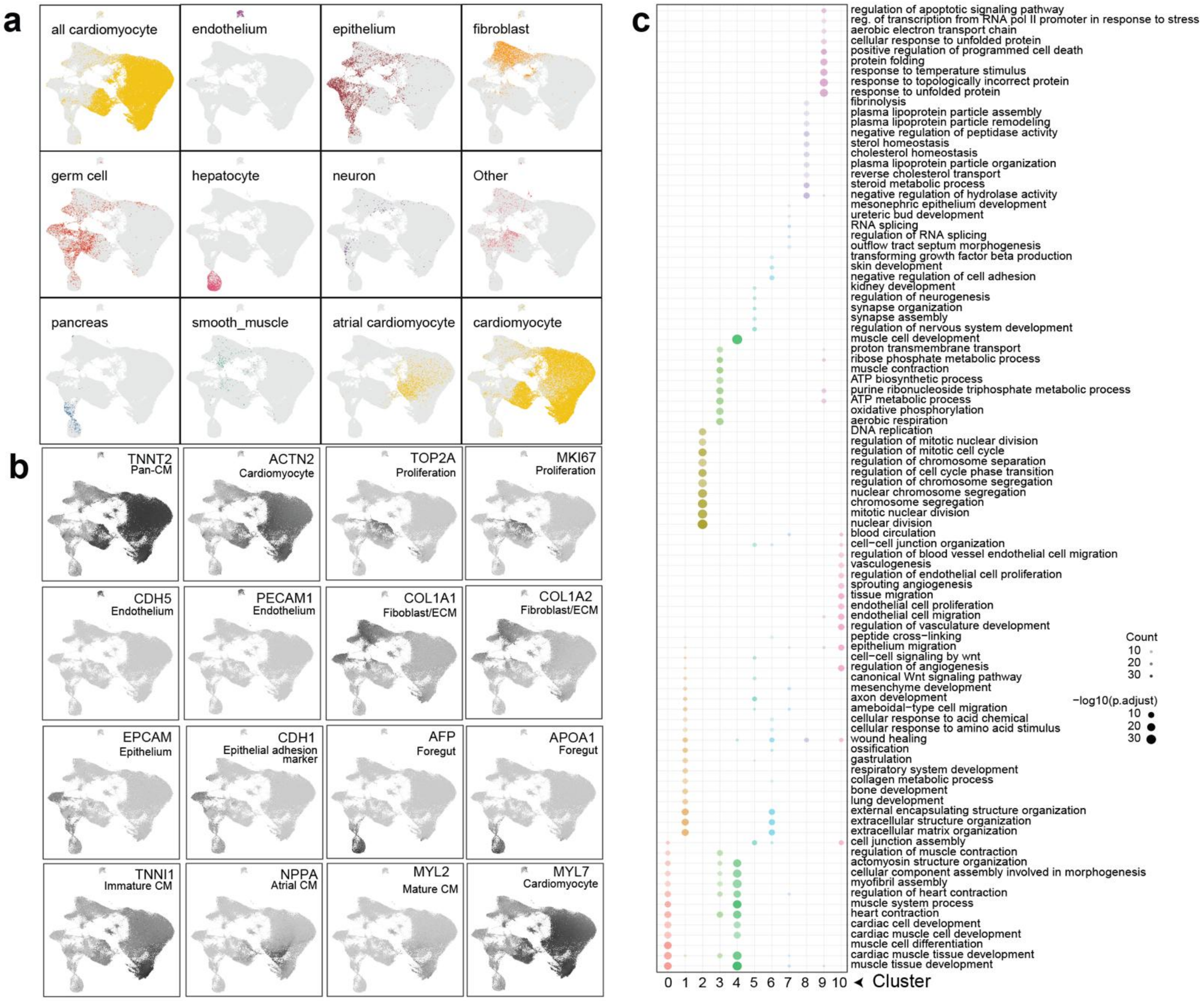


**Figure S2 | Supporting analyses for cell type identification in integrated single-nucleus RNA-seq datasets.**

**a,** scGPT-based cell type annotation across the integrated single-nucleus RNA-seq dataset. UMAP projections display predicted cell identities including cardiomyocytes, atrial cardiomyocytes, endothelium, fibroblasts, epithelium, smooth muscle, foregut endoderm, germ cell–like populations and additional minor cell types. Highlighted populations correspond to predicted classifications assigned by the scGPT annotation model.

**b,** Expression of canonical marker genes used to validate cluster annotations. UMAP projections show representative markers for cardiomyocytes (TNNT2, ACTN2), proliferative cells (TOP2A, MKI67), endothelial cells (CDH5, PECAM1), fibroblast/extracellular matrix populations (COL1A1, COL1A2), epithelial cells (EPCAM, CDH1), foregut/endodermal populations (AFP, APOA1), and cardiomyocyte subtype markers including immature cardiomyocytes (TNNI1), atrial cardiomyocytes (NPPA) and mature cardiomyocyte markers (MYL2, MYL7).

**c,** Gene Ontology enrichment analysis of cluster-specific gene expression programs across the integrated dataset. Dot plot shows representative biological processes enriched among the top differentially expressed genes for each cluster, with dot size representing the number of genes contributing to the enrichment and colour indicating adjusted significance (−log10 adjusted P value). Enriched terms support annotation of clusters corresponding to cardiomyocyte differentiation and muscle contraction, cell cycle and proliferation, vascular development, epithelial and mesenchymal processes, extracellular matrix organization and metabolic processes.

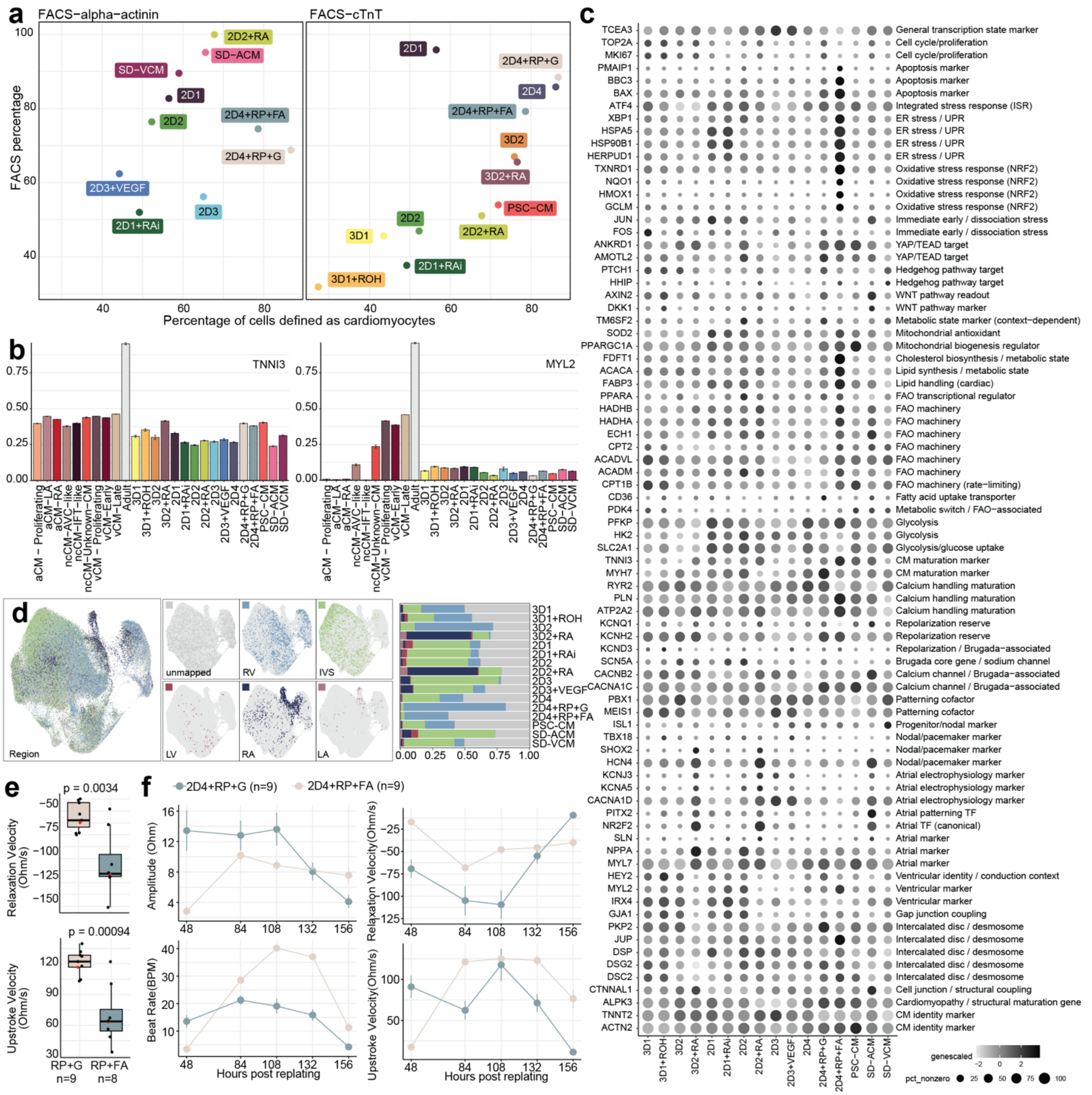


**Figure S3 | Supporting analyses for cardiomyocyte subtype classification and maturation states across differentiation protocols.**

**a,** Flow cytometry validation of cardiomyocyte yield across differentiation protocols. Scatter plots show the relationship between the percentage of cells classified as cardiomyocytes by single-nucleus RNA sequencing and the percentage of cardiomyocytes measured by flow cytometry using **α-actinin** (left) or **cardiac troponin T (cTnT)** (right) staining.

**b,** Expression of cardiomyocyte subtype markers across clusters and differentiation protocols. Bar plots show relative expression of representative markers including **TNNI3** (ventricular/mature cardiomyocyte marker) and **MYL2** (ventricular cardiomyocyte marker), highlighting variation in cardiomyocyte subtype composition across differentiation conditions.

**c,** Expression of lineage, stress-response and maturation gene programs across differentiation protocols. Dot plot shows scaled expression of selected genes representing transcriptional state markers including proliferation and cell-cycle genes, apoptotic and stress-response pathways (including ER stress and oxidative stress programs), metabolic regulators, fatty acid oxidation machinery, glycolytic and mitochondrial pathways, electrophysiological markers and chamber-specific cardiomyocyte identity genes. Dot size represents the proportion of cells expressing each gene and colour indicates scaled expression level.

**d,** Regional annotation of cardiomyocyte identities based on label transfer to a human fetal heart reference atlas. UMAP projections highlight predicted anatomical regions including right ventricle (RV), left ventricle (LV), right atrium (RA), left atrium (LA) and interventricular septum (IVS). Stacked bar plots summarize the proportion of region-specific cardiomyocyte identities across differentiation protocols.

**e,** Functional comparison of cardiomyocyte maturation between replated (2D4+RP+G) and fatty acid–matured (2D4+RP+FA) cardiomyocytes. Box plots show differences in relaxation velocity and upstroke velocity derived from contractility measurements.

**f,** Time-course analysis of cardiomyocyte contractility following replating or metabolic maturation. Line plots show changes in contractile amplitude, beat rate, relaxation velocity and upstroke velocity across time following replating in the two maturation conditions (2D4+RP+G and 2D4+RP+FA), demonstrating progressive functional divergence associated with metabolic maturation.

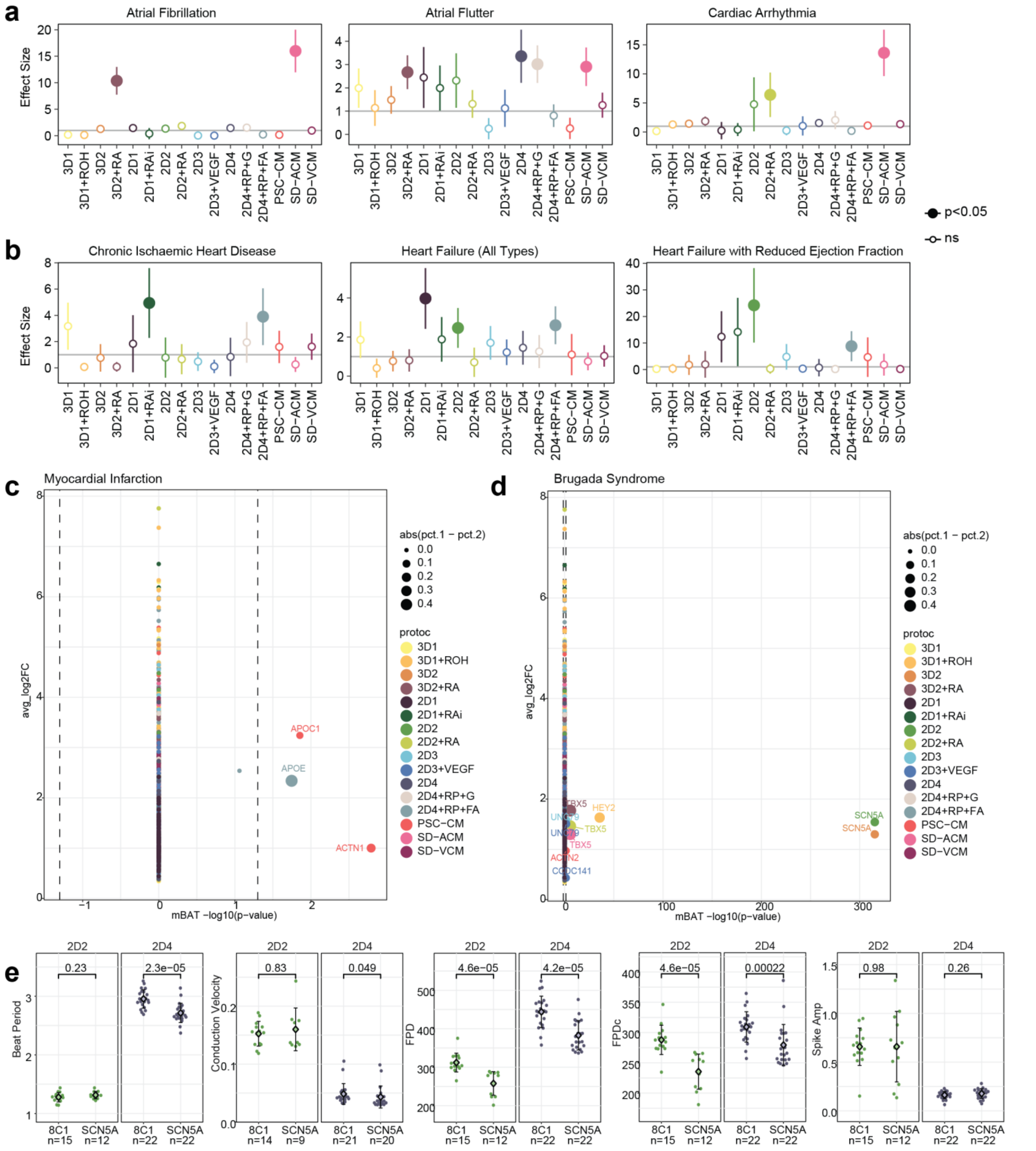


**Figure S4 | Additional analyses supporting protocol–disease associations and electrophysiological disease modelling.**

**a,** Enrichment of atrial arrhythmia–related genetic architectures across differentiation protocols. Effect sizes derived from SBayesRC analyses are shown for atrial fibrillation, atrial flutter and cardiac arrhythmia GWAS traits. Points represent protocol-specific enrichment estimates with error bars indicating uncertainty. Filled points denote statistically significant associations ($P < 0.05$).

**b,** Enrichment of genetic architectures associated with ischemic and heart failure–related traits across differentiation protocols. Effect sizes are shown for chronic ischemic heart disease, heart failure (all types) and heart failure with reduced ejection fraction. Protocol-dependent differences highlight variation in disease-relevant transcriptional programs across cardiomyocyte states.

**c,** Gene-level association analysis linking myocardial infarction GWAS signals to protocol-specific gene expression programs. Scatter plot shows average gene expression ($\log_2$ fold change) against gene-level association significance (mBAT $-\log_{10}P$). Highlighted genes include **APOC1**, **APOE**, and **ACTN1**, illustrating enrichment of lipid metabolism–related loci within fatty acid–matured cardiomyocyte states.

**d,** Gene-level association analysis linking Brugada syndrome GWAS signals to protocol-specific transcriptional programs. Scatter plot highlights enrichment of sodium channel and cardiac conduction–related genes including **SCN5A**, **SCN10A**, **TBX5** and **HEY2**, supporting the association between ventricular differentiation protocols and Brugada syndrome genetic architecture.

**e,** Additional electrophysiological measurements in control (8C1) and **SCN5A** mutant cardiomyocytes differentiated using ventricular protocols. Box plots show beat period, conduction velocity, field potential duration (FPD), corrected field potential duration (FPDc) and spike amplitude across protocols (2D2 and 2D4). Statistical comparisons between control and mutant lines are indicated above each panel. These data demonstrate protocol-dependent sensitivity for detecting electrophysiological phenotypes associated with SCN5A-mediated channelopathy.
